## Supplemental Table 7 for "Regulation of heme utilization and homeostasis in *Candida albicans*"

**Table 3: List of primers**

| # | Name | Sequence |
|---|---|---|
| **1** | F1-Transposon-FPNI | CCGTTCGTTTTCGTTACCGGTATATC |
| **2** | F2-Transposon-FPNI | CCGTCCCGCAAGTTAAATATG |
| **3** | F3-Transposon-FPNI | GTATTTTACCGACCGTTACCGACC |
| **4** | R1-FPNI | CGCAGGTGACATAGATGC |
| **5** | R2-FPNI | GACATAGATGCTTAGCGCTGAGG |
| **6** | R3-FPNI | CGCAGGTGACATAGATGCTTAGCG |
| **7** | PstI-HUF1 (+1) | GGCTGCAGATGGGTCCTATAGCTGTAA |
| **8** | HUF1-HindIII (+3255) | CCAAGCTTTCCTTGAAAGTAGTTTTCTATATT |
| **9** | SpeI- HUF1 (-1890) | GCGGCCGCTCTAGAACTAGTTCTAATAGCTTCCCACGG TATG |
| **10** | HUF1-HindIII (+3636) | TCGACGGTATCGATAAGCTTCTCAATATACTAGACCCT GG |
| **11** | HUF1-1 (-491 fwd) | GGATGTTATTTAGAATCAC |
| **12** | HUF1-3 (+24 rev) | cacggcgcgcctagcagcggGAAACTATTACTGCTTTCC |
| **13** | HUF1-4 (+3243 fwd) | gtcagcggccgcatccctgcCTACTTTCAAGGATAAATTTG |
| **14** | HUF1-6 (+3713 rev) | GTTCGATTTATAGGAAACAG |
| **15** | SpeI-HUF1 (+1258) | gcgactAGTTGAAAATCAGTGGGATTTA |
| **16** | HUF1-XhoI (+3255) | cgctcgagTCCTTGAAAGTAGTTTTCTATAT |
| **15** | ACT1f | TGAAGCCCAATCCAAAAGAGG |
| **16** | ACT1r | TTTCCATATCGTCCCAGTTGG |
| **17** | FRP2f | CGACAAGGTCAAAACAATGCG |
| **18** | FRP2r | GAGAAATGCCACTGAGTCAAATG |

| | | |
|---|---|---|
| **19** | HUF1 60 fwd | ACCTAACAAAGTCCAGAAACCT |
| **20** | HUF1 145 rev | CGCATGCTGGTTTACCTTTATC |
| **21** | HMX1 fwd | GCACGATAGAGCAGACAAGACAG |
| **22** | HMX1 rev | TCCGGTTTCCAAAACTTGCTCC |
| **23** | PGA7 FWD | TTCCTCGATGCTTTCCACTGCC |
| **24** | PGA7 REV | ATGAGCCTGTAGATGACGAGCC |
| **25** | FRP1 FWD | GGAAACAAAGGAAGAATTGCCAC |
| **26** | FRP1 rev | CACCTTTCACCCTTCTAGCAC |
| **27** | RBT5f | TGCTCGCCTTATCCTTATTGTC |
| **28** | RBT5r | GTTGATGGAAGCGGTTTTAGC |
