## Supplemental figures for "Regulation of heme utilization and homeostasis in *Candida albicans*"

### SUPPLEMENTARY FIGURES
#### Andrawes et al.

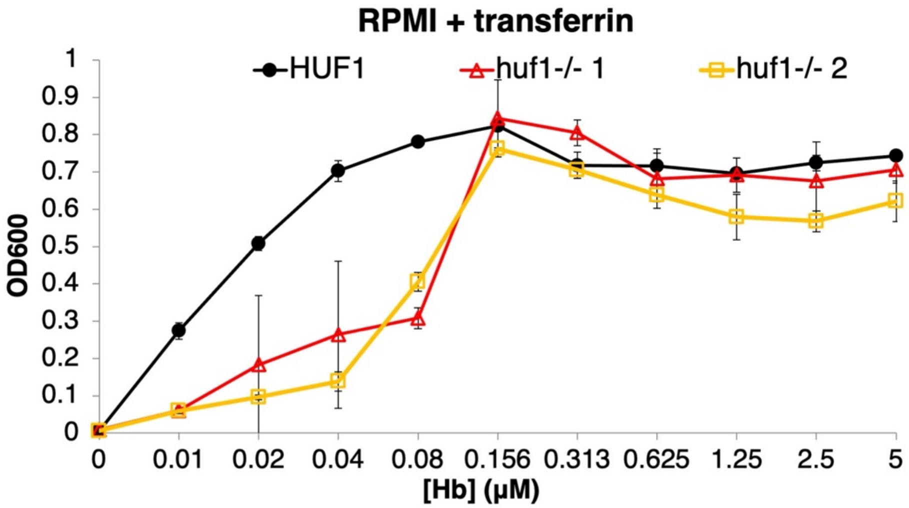


**Fig. S1.** The *huf1*$^{-/-}$ mutant is defective in growth on hemoglobin in RPMI medium supplemented with transferrin. Two independent *huf1*$^{-/-}$ mutant clones from the Homann collection [14], together with wild-type strain SN148, were inoculated in desferrated RPMI medium supplemented with transferrin and with increasing amounts of hemoglobin, as indicated. The wild-type and mutant strains were inoculated in triplicate in 96 well plates, and incubated for 2 days at 30°C. The graph indicates the average density for each triplicate, and the error bars indicate the standard deviations.

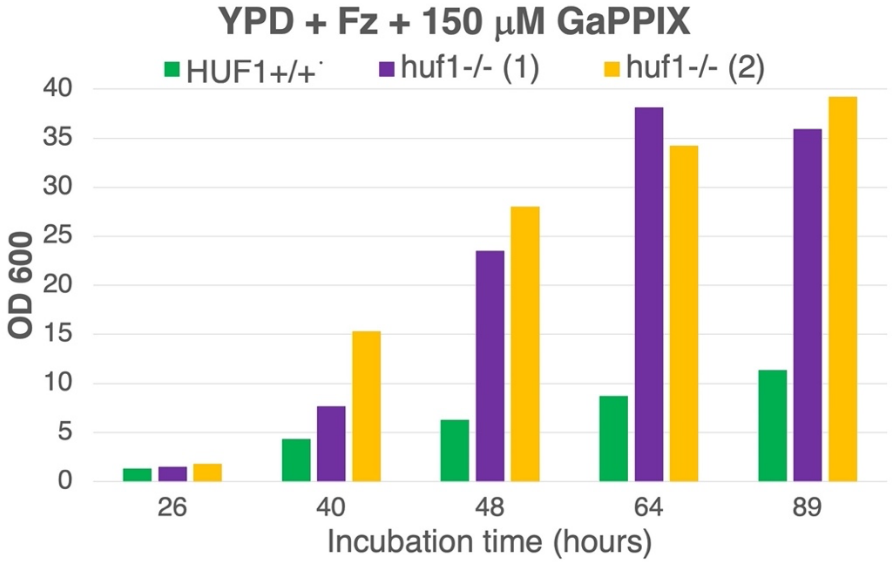


**Fig. S2.** The *huf1*$^{-/-}$ mutant grows faster in the presence of GaPPIX. The wild-type strain SN148 and two independent *huf1*$^{-/-}$ mutant strains (KC1406) were inoculated at OD600 = 0.0005 in YPD supplemented with 1 mM ferrozine and 150 μM GaPPIX, and incubated at 30°C in 50 ml Falcon tubes with vigorous shaking. The densities were measured at the indicated times after inoculation.

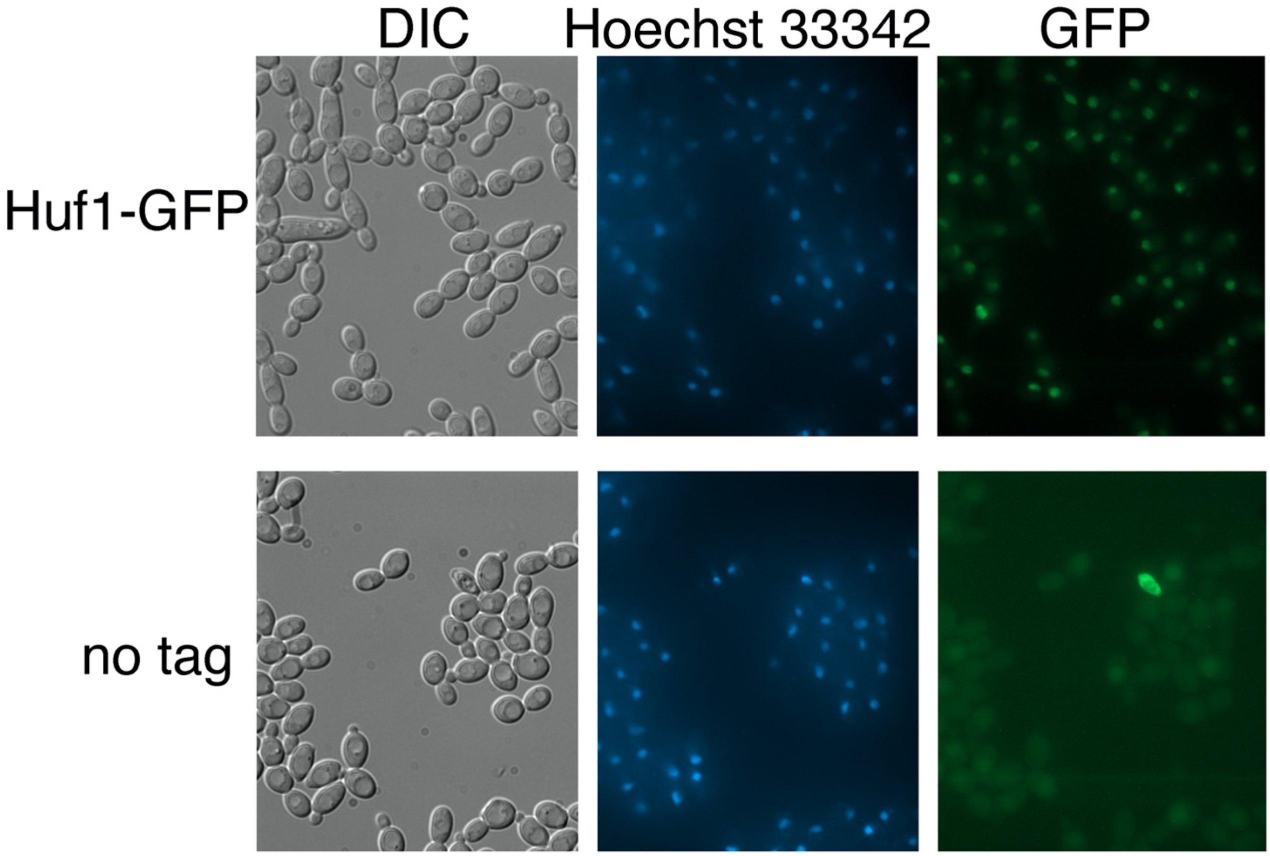


**Fig. S3.** Huf1-GFP is located in the nucleus. The strain expressing Huf1-GFP under its native promoter, and a control untagged strain, were grown 4 h in YPD medium containing 1mM ferrozine and 10 µM hemin, then briefly stained with the DNA stain Hoechst 33342 and visualized by epifluorescence microscopy.
